# Modulation of *Saprolegnia parasitica* growth with copper and ionophores

**DOI:** 10.1101/2024.06.17.599375

**Authors:** Tomisin Happy Ogunwa, Madison Grace Thornhill, Daniel Ledezma, Ryan Loren Peterson

## Abstract

*Saprolegnia parasitica* is an oomycete pathogen responsible for saprolegniasis diseases that result in large production losses in the catfish and salmon aquaculture industry. The use of copper sulfate as an anti-Saprolegnia treatment has been reported as an alternative to malachite green, formaldehyde and hydrogen peroxide treatment methods. The current study investigates a new strategy to inhibit *Saprolegnia parasitica* growth by combining copper and ionophores at low levels. The chemical agents tetraethylthiuram disulfide (TDD), ciclopirox olamine (CLP), 2-mercaptopyridine N-oxide (MPO), 5-chloro-8-hydroxy-7-iodoquinoline (CHI), 5,7-dichloro-8-hydroxyquinoline (DHQ) and 8-Quinolinol (8QN) were identified to inhibit *S. parasitica* growth in a copper-dependent manner. At concentrations below the lethal dose of individual ionophore, increasing copper concentrations resulted in synergetic *S. parasitica* growth inhibition. The addition of the exogenous copper chelator bathocuproine sulfate (BCS), reversed the inhibition of *S. parasitica* growth by TDD, CLP, MPO, and 8QN but not CHI and DHQ. Our data demonstrates that ionophores, in combination with low levels of copper, can effectively limit *S. parasitica* growth both in a liquid and solid support growth environment. Investigations into the underlying mechanism of Cu-ionophore toxicity are discussed.

## 1. INTRODUCTION

*Saprolegnia parasitica* is a devastating oomycete pathogen infecting salmon, trout, catfish, and ornamental fish species commonly found as a white cotton-like growths on wounded and vulnerable adult fish as well as developing embryos (Bruno et al., 2011). Saprolegniasis is prevalent in hatcheries, but environmental factors, including poor water quality, rapid temperature changes, and wounds, can increase animal susceptibility to infection. Saprolegniasis leads to severe yield losses (∼ 10 percent loss) in fish production with a concomitant reduction in the quality of fish and farmers’ profits (Earle & Hintz, 2014). Despite the various efforts applied to control the disease over the years, very little progress has been made towards developing new United States Food and Drug Administration (FDA)-approved methods for Saprolegniasis treatment and prevention, notwithstanding the significant efforts to approve Cu-sulfate (Straus et al., 2016; Straus et al., 2020; Straus, Mitchell, Radomski, et al., 2009) and peracetic acid (Good et al., 2020) as chemical treatment methods. Currently, only two products are FDA approved for saprolegnia treatment which include PEROX-AID and Parasite-S (Haskell et al., 2004).

Malachite green, hydrogen peroxide and formaldehyde are among the agents previously adopted to combat saprolegniasis in fish production (Gaikowski et al., 1998; Stammati et al., 2005; Walser & Phelps, 1994). Although some of these fungicides proved beneficial in the control and prevention of the disease, malachite green is now banned worldwide due to environmental impact, including teratogenic and carcinogenic effects on exposed rainbow trout, egg and fry (Srivastava et al., 2004). Newer alternative saprolegniasis chemical treatment methods have included the use of Cu-sulfate, peractic acid, Clotrimazole, and Saprolmycin A-E (Nakagawa et al., 2012; Warrilow et al., 2014). Additionally, natural products have been tested and shown to control *S. parasitica* growth. Among these plants, *Punica granatum, Cyperus esculentus*, *Carthamus tinctorius* and *Thymus vulgaris* showed potential to suppress mycelial growth of *Saprolegnia diclina in vitro* at various concentrations (Emara et al., 2020; Mostafa et al., 2020; Xue-Gang et al., 2013). However, the mechanism and chemical compounds resulting in anti-oomycete activity are not well defined (Caruana et al., 2012).

There has been considerable effort to approve copper sulfate as a cost-effective treatment method to control *S. parasitica* infection in catfish (Straus, Mitchell, Carter, et al., 2009; Straus et al., 2011; Sun et al., 2014). One drawback is that local water quality and hardness can impact copper solubility and overall treatment effectiveness (Straus, Mitchell, Carter, et al., 2009). We hypothesized that small copper-binding agents, such as the general class of ionophores, may overcome the copper solubility challenges experienced with local water quality while simultaneously assisting in the delivery of copper ions into the pathogen cytoplasm. Herein, we report a new strategy designed to inhibit *S. parasitica* growth, which combines ionophore small molecules with copper sulfate. The data obtained in our experiments suggest that copper can modulate the growth of *S. parasitica* when combined with selected ionophores, even at minimal concentrations. This opens new potential anti-oomycete therapeutic avenues that may be more efficient than employing Cu-sulfate alone to combat outbreaks in an aquaculture setting.

## 2. MATERIALS AND METHODS

### 2.1 Chemicals

All chemical agents used in this study were purchased from commercial vendors at >98% purity levels and used as received. Milli-Q (18Ω-resistance) water was used for the preparation of all media and solutions. Chemical compounds including bathocuproine sulfate (BCS), tetraethylthiuram disulfide (TDD), ciclopirox olamine (CLP), 2-mercaptopyridine N-oxide (MPO), 5-chloro-8-hydroxy-7-iodoquinoline (CHI), 5,7-dichloro-8-hydroxyquinoline (DHQ), 8-Quinolinol (8QN), bicinchonic acid disodium salt (BCA), 2-thiophene carboxylic acid (TCA), ammonium tetrathiomolybdate (ATM), quercetin (QUE), curcumin (CUR), azelaic acid (AZE), copper (II) dibutyldithiocarbamate (CDT), bathophenanthroline disulfonic acid disodium salt (BPS), 2-mercaptopyridine N-oxide zinc salt (MPOZ), kaempferol (KAE) and ethylenedinitrilotetraacetic acid disodium salt dihydrate (Na-EDTA) were purchased from vendors in the United States.

### 2.3 Preparation of 2222X growth media

Chemically defined 2222X growth media was developed for this project based on growth experiments described by Powell et al. and consisted of 2 g/L monosodium glutamate, 2 g/L glucose, 3.4 g/L yeast nitrogen base (YNB), and 200 mg/L methionine at a pH of 6.0 and sterilized via ultra-filtration (Powell et al., 1972). Solid 2222X plates were made using 15 g/L agar added prior to autoclaving.

### 2.3 Maintenance and propagation of *Saprolegnia parasitica*

*Saprolegnia parasitica* CBS223.65 reference strain, purchased from the Westerdijk Fungal Biodiversity Institute (Utrecht, Netherlands), was used for all experiments in this study and maintained on the 2222X agar plates. Transfer of *S. parasitica* samples from plate to plate or experimental tubes was done using disposable sterile 3 mm biopsy punches (Robbins Instruments, India).

### 2.4 *In vitro* screening protocol

A 3 mm agar plug of *S. parasitica* was used to inoculate a 10 ml of 2222X solution in a 50 ml vented conical tube and incubated at 28 °C with 220 rpm for 5 days. The off-white *S. parasitica* filamentous mycelium was removed with forceps, drained, and transferred to a pre-weighed spin filter tube (CLS9301), and centrifuged for 2 minutes at 8000 X g to remove excess liquid prior to weighing.

For the initial screening of metal-binding compounds against *S. parasitica* growth, each chemical compound (10 μM final concentration using a 50mM stock in DMSO) was added to a culture tube containing 10 ml of 2222X media supplemented with 100 μM Cu-sulfate and cultured as described above. Only compounds exhibiting total growth inhibition were further evaluated. For IC_50_ determination, the concentration of each metal-binding compound was varied as appropriate: TDD (0 – 50 μM), CLP (0 – 2 μM), MPO (0 – 2 μM), CHI (0 – 50 μM), DHQ (0 – 50 μM), and 8QN (0 – 10 μM) in a 10 ml 2222X media and cultured as described above.

### 2.5 Oxyblot determination of oxidized and carbonylated proteins

To investigate the ability of copper and ionophore mixtures to induce metal-mediated oxidative stress in *S. parasitica,* we quantified protein oxidation levels in *S. parasitica* lysates using the OxyBlot Protein Oxidation Detection kit (Millipore, Billerica, MA, USA). Carbonylated proteins were derivatized with 2,4 Dinitrophenyl hydrazine (DNP) reagent and assayed via western blot using an anti-DNP primary antibody as described below. In brief, *S. parasitica* samples were grown for four days and then acutely exposed to various treatments including copper, BCS and MPO for two hours. The samples were harvested and lysed in the presence of 50 mM DTT and proteinase inhibitors. Each lysate was centrifuged at 17,000g for 10 mins at 4°C and the supernatant was used in the derivatization step. The carbonyl groups in *S. parasitica* protein samples (80 μg) were derivatized to 2,4-dinitrophenylhydrazone (DNP) via a reaction with 2,4-dinitrophenylhydrazine (DNPH) in the presence of 6% Sodium dodecyl sulfate (SDS). After incubation at 25°C for 15 minutes, the reaction was quenched by adding the neutralization solution. Next, 4-12% SDS-PAGE was carried out on the derivatized samples. The samples were then transferred to a nitrocellulose membrane and incubated with rabbit anti-DNP (1:150 dilution) for 1hr at RT. Goat anti-rabbit Alexa fluor 647 (1:300 dilution) was added, and the blot developed according to western blotting protocol. Finally, images were generated using the Image Lab software (Bio-Rad).

### 2.6 Data analysis

The growth inhibition of *S. parasitica* with selected ionophores were analyzed using Microsoft excel package and presented as the mean percentage of growth compared to control conditions (i.e. 2222X media alone). Images were taken with a digital camera.

## 3. RESULTS

The inoculation of liquid 2222X growth media with a 4 mm gel plug of *S. parasitica,* cultured on 2222X plates, leads to the reproducible culture of *S. parasitica* which can be achieved in a standard shaking incubator at 28 °C (Fig. 1). Using this protocol, we determined the lethal dose of Cu^2+^, as CuSO_4_, against *S. parasitica in vitro*. The IC_50_ value for Cu^2+^ was ∼100 μM while a concentration of 200 μM Cu^2+^ completely inhibited *S. parasitica* growth (Fig. 2). To identify chemical compounds that could enhance Cu^2+^ toxicity against *S. parasitica*, we carried out a screening experiment on nineteen chemical compounds at a concentration of 10 μM with and without supplemental 100 μM Cu^2+^. These experiments revealed six compounds with copper-dependent activity against the *S. parasitica* growth (Fig. 3). Tetraethylthiuram disulfide (TDD), ciclopirox olamine (CLP), 2-mercaptopyridine N-oxide (MPO) and 8-Quinolinol (8QN), chemical compounds highlighted in green, inhibited *S. parasitica* growth in copper-dependent manner whereas those in blue including 5-chloro-8-hydroxy-7-iodoquinoline (CHI) and 5,7-dichloro-8-hydroxyquinoline (DHQ) displayed copper-independent growth inhibition. In addition, compounds highlighted in gray moderately suppressed *S. parasitica* growth at 100 μM concentration. However, those shaded in white demonstrated negligible effects on *S. parasitica* growth. Chemical compounds highlighted in green were selected for further characterization as they exhibited copper-dependent activity (vide infra).

**Fig. 1.**
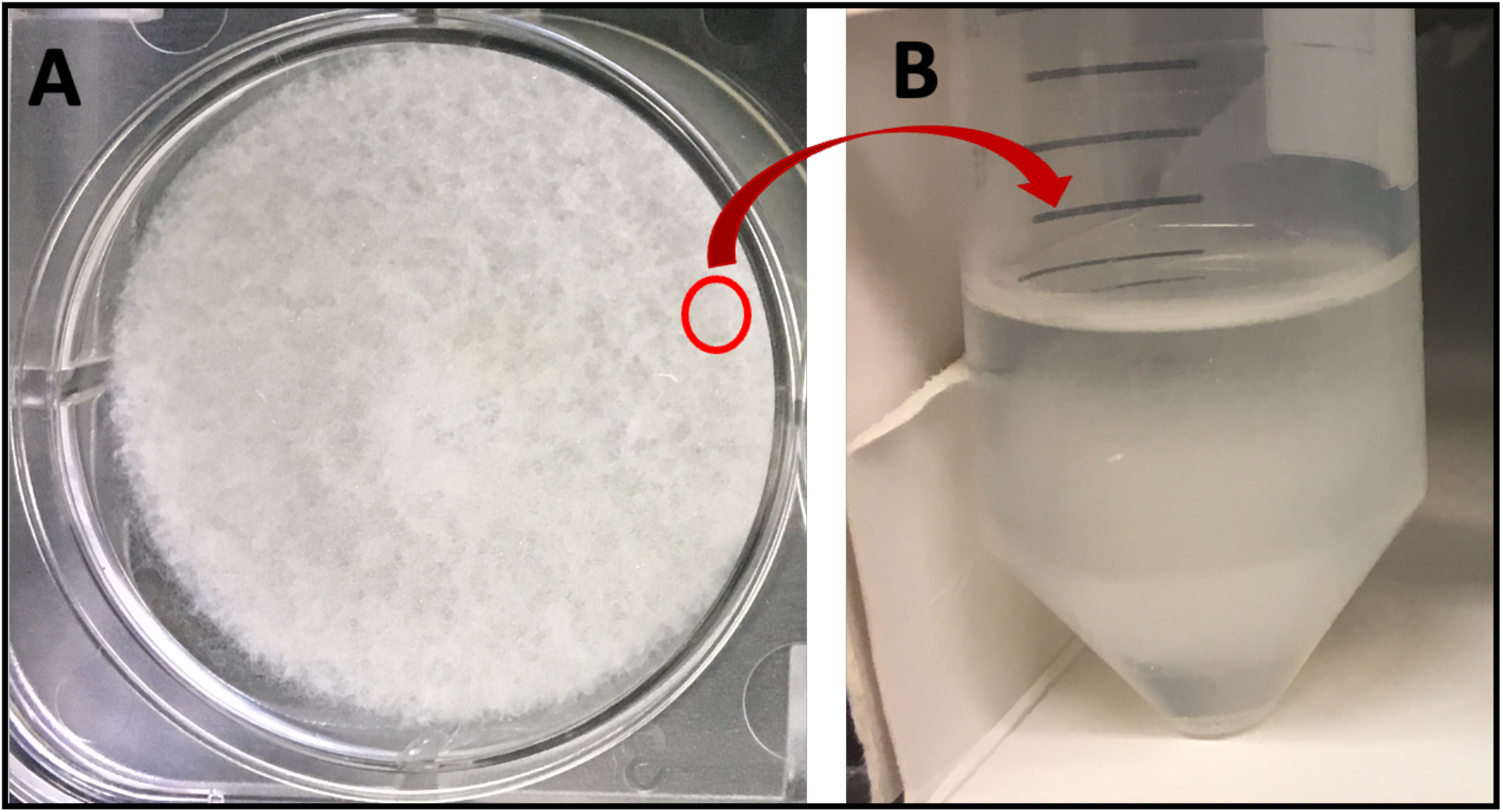
*Saprolegnia parasitica* growth behavior on synthetic 2222X growth media: (a) solid agar plate cultured for 5 days at room temperature. (b) *S. parasitica* cultured from a 4 mm gel punch and cultured in 10mL of 2222X media at 28°C and 220 rpm for 5 days.

**Fig. 2.**
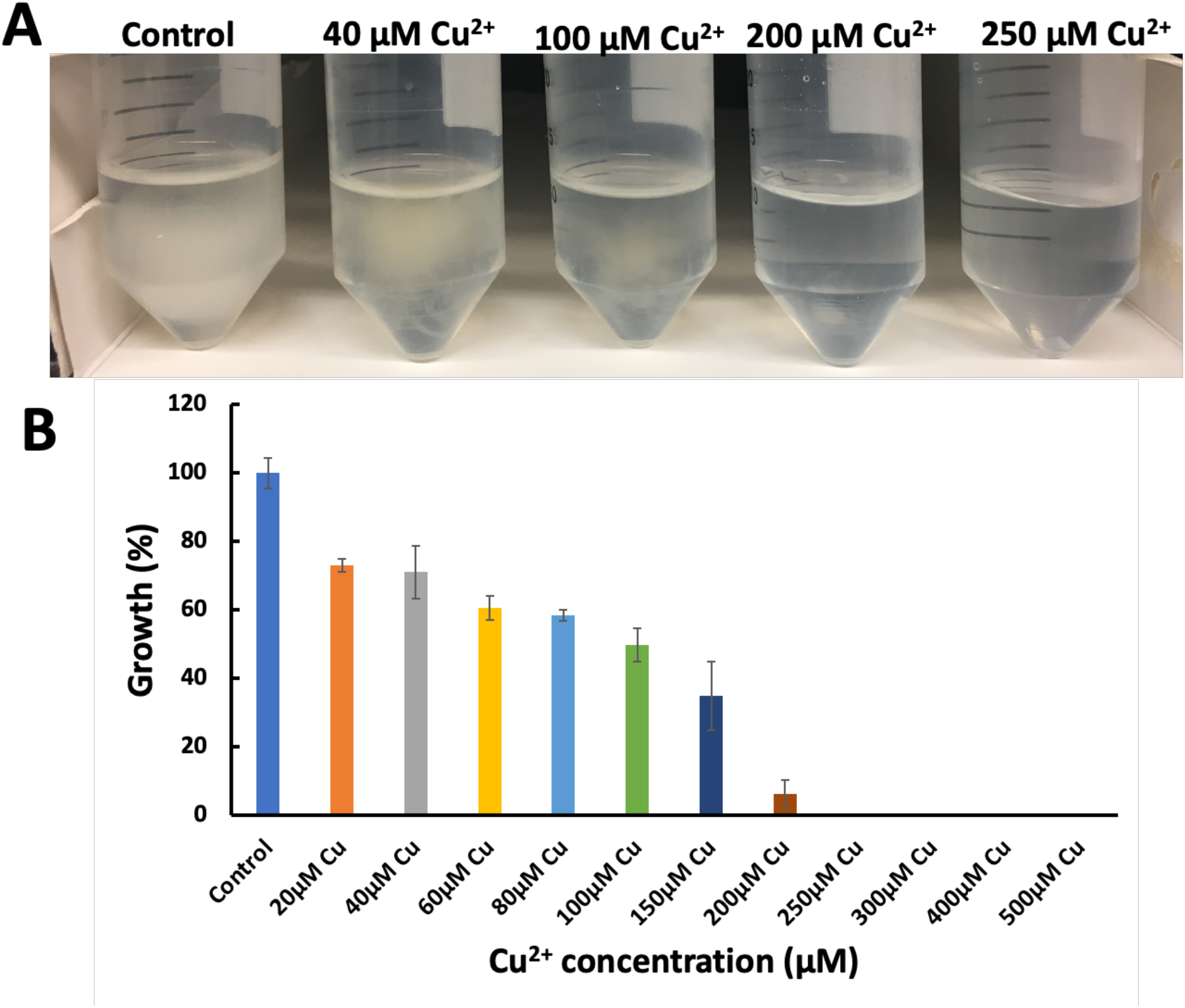
Effect of copper (Cu^2+^) on the growth of *S. parasitica* in synthetically defined 2222X media. Total growth was measured in 2222X media after 5-day incubation at 28 °C and 222 rpm. **A.** Addition of Cu^2+^ induced a brownish color in *S. parasitica* compared to the control. **B.** The growth behavior of *S. parasitica* as a function of supplemental copper concentration. An IC_50_ for Cu^2+^ was 100 μM, whereas no growth was detected at a copper concentration > 200 μM. Error bars represent the standard deviation of the mean of data from triplicate experiments.

**Fig. 3.**
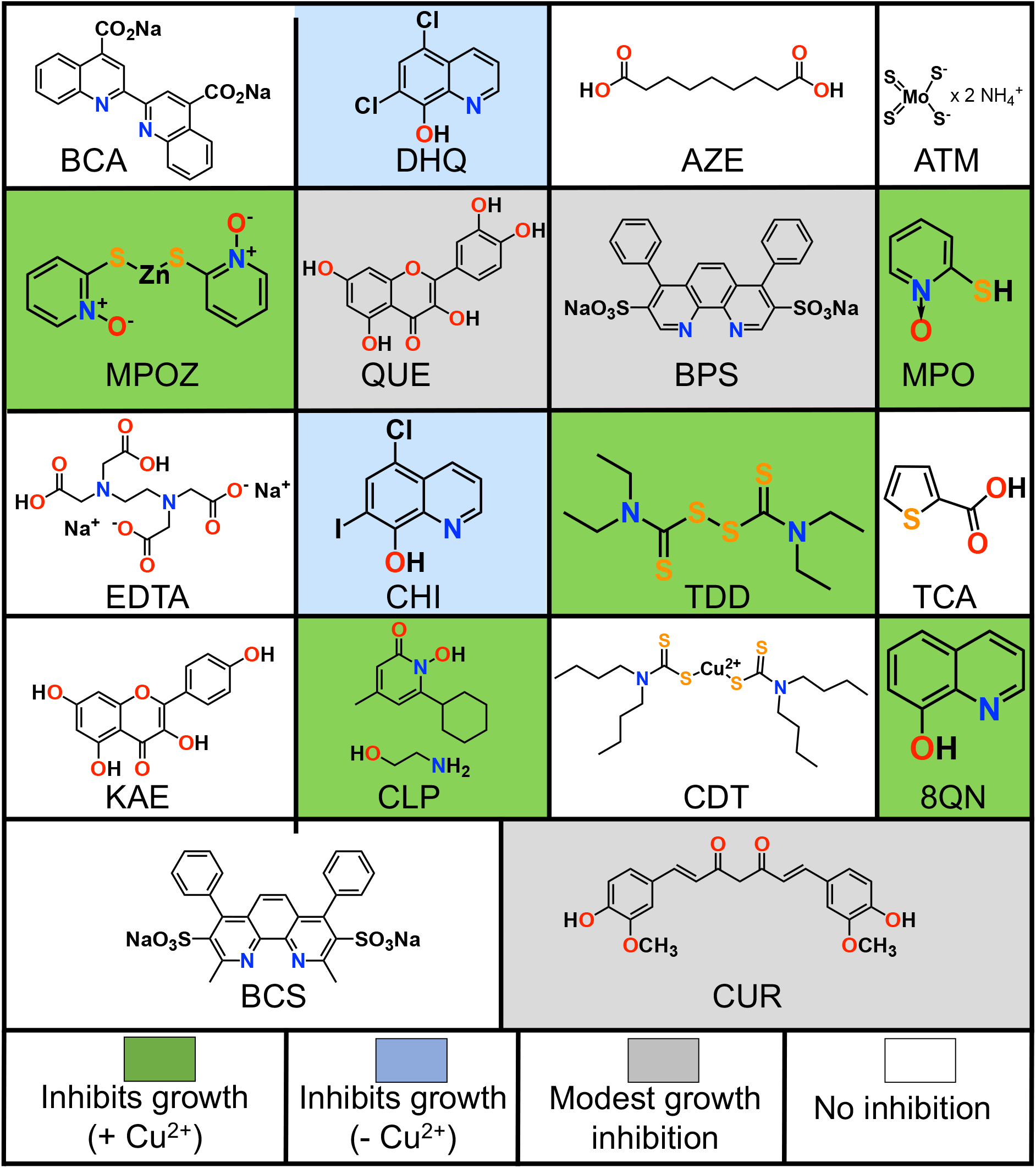
Chemical structure and metal-dependent activities of select chemical compounds used in this study. *In vitro* screening was achieved using doses of 10 μM of each chemical compound in the presence or absence of supplemental CuSO_4_ (up to 100 μM) in 2222X media. Compounds highlighted in green inhibited *S. parasitica* growth with added Cu^2+^ whereas compounds shaded blue exhibited growth inhibition without Cu^2+^ dependency. Compounds highlighted as gray showed only moderate effect. However, those that displayed negligible effect on *S. parasitica* growth *in vitro* are highlighted in white (Helsel et al., 2017; Tedesco et al., 2019).

### 3.1 Dose-dependent effect of ionophores on *S. parasitica* growth *in vitro*

To establish the lethal dose for each of the ionophores against *S. parasitica*, a concentration-dependent experiment was carried out on the selected (six) copper-dependent ionophores. These experiments revealed a range in IC_50_ values from < 0.6 to >15 μM (Figure 4). The least effective compound was TDD with an estimated 17 μM concentration needed to suppress ∼ 50% to *S. parasitica* growth (Fig. 4a). CLP and MPO exhibited high toxicity with a lethal dose of 1 μM each (Fig. 4b, c) whereas CHI caused *S. parasitica* death at approximately 4 μM (Fig. 4d). DHQ and 8QN were the most effective compounds at preventing *S. parasitica* growth with IC_50_ concentration of approximately 0.5 μM (Fig. 4e, F). However, 8QN required a much higher concentration of 8 μM to completely inhibit *S. parasitica* propagation. Notably, all the selected ionophores showed a concentration-dependent pattern in suppressing *S. parasitica* growth *in vitro*.

**Fig. 4.**
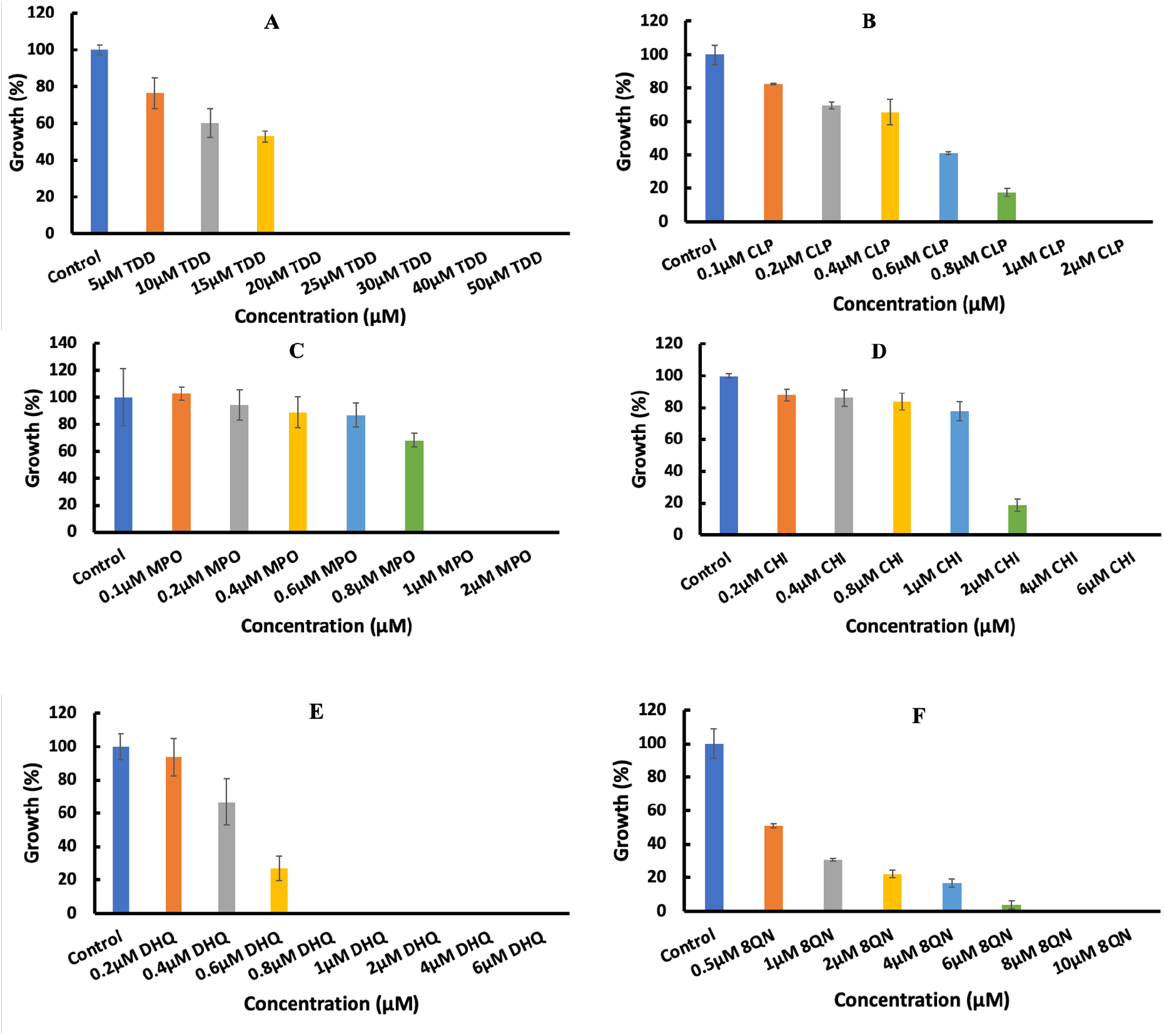
Dose-dependent assay of ionophores against *S. parasitica*. Lethal dose for each ionophore was determined *in vitro* as (a) TDD (20 μM), (b) CLP (1 μM), (c) MPO (1 μM), (d) CHI (4 μM), (e) DHQ (0.8 μM) and (f) 8QN (0.5 μM). Growth was measured at 28°C after 5 days and normalized to the growth of control. Error bars represent the standard deviation of the mean of data from triplicate experiments.

### 3.2 Effect of copper-ionophore on *S. parasitica* growth

To determine whether the selected ionophores can boost Cu^2+^ toxicity against *S. parasitica*, we varied the copper concentration at a fixed ionophore dose. A unique concentration for each ionophore below the IC_50_ was selected. Interestingly, a combination of 10 μM TDD with 16 μM copper completely inhibited *S. parasitica* (Fig. 5a) growth. For CLP, 0.6 μM of the ionophore with copper led to the dose-dependent suppression of *S. parasitica* growth (Fig. 5b). Increasing the dose of CLP to 0.8 μM yielded a total inhibition of the oomycete growth at 4 μM copper (Fig. S1). MPO exhibited a high sensitivity to copper by killing *S. parasitica* at a dose of 0.4 μM and 1 μM, respectively (Fig. 5c). Using a higher MPO dose (0.6 μM) resulted in the total death of *S. parasitica* at 0.5 μM copper (Fig. S2). However, lowering the MPO concentration to 0.2 μM was not completely lethal to the pathogen in the presence of 100 μM copper (Fig. S3). These results are consistent with the data on MPZO, the zinc salt of the pyrithione (MPO) which exhibited reversible Cu^2+^-dependent toxicity against *S. parasitica* (Fig. S4).

**Fig. 5.**
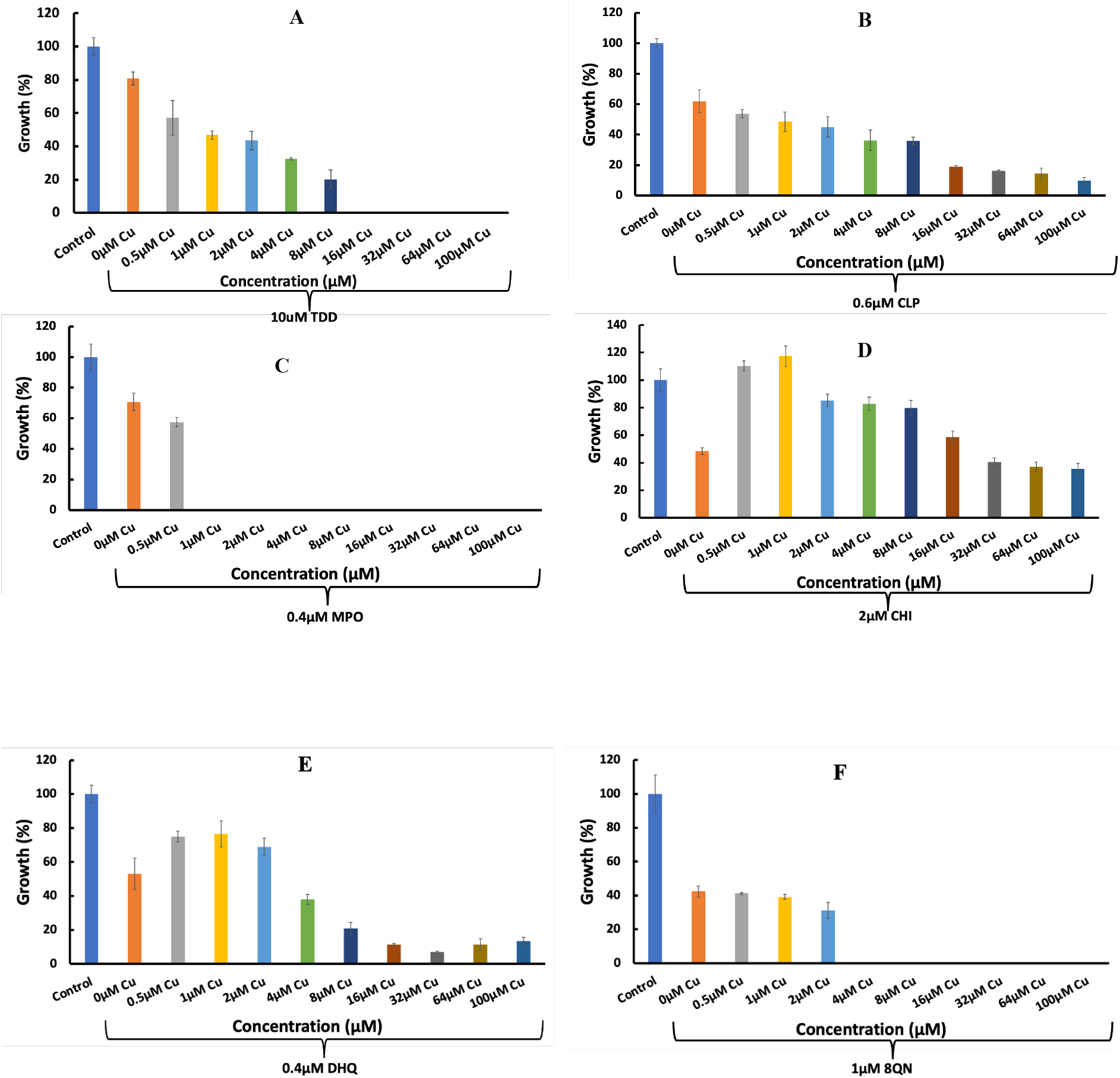
Effect of copper - ionophore on *S. parasitica* growth *in vitro*. A low dose for each ionophore (a) 10 μM TDD, (b) 1 μM CLP, (c) 1 μM MPO, (d) 4 μM CHI, (e) 0.8 μM DHQ and (f) 8 μM 8QN was combined with increasing concentration of Cu^2+^ to determine the precise Cu^2+^-ionophore dose that can potently suppress *S. parasitica* growth. Growth was measured at 28°C after 5 days and normalized to the growth of control. Error bars represent the standard deviation of the mean of data from triplicate experiments.

For the chemicals CHI and DHQ, a sigmoidal copper dose response is observed (Fig. 5d, e). Specifically, at 1 μM CHI and 0.4 μM DHQ, a low level of copper appeared to boost the growth of *S. parasitica* up to 2 μM copper (Fig. 5d, e). Repeating the experiment at a higher CHI dose (2 μM) yielded a consistent effect showing a sigmoidal curve on *S. parasitica* growth (Fig. S5). Similarly, the concentration of DHQ was increased in the presence of 100 μM copper and the results corroborated previous observation that *S. parasitica* can survive a combined 100 μM Cu^2+^ and 0.6 μM DHQ (Fig. S6). However, the ionophore 8QN with a relatively lesser toxicity against *S. parasitica* in the earlier dose-dependent assay demonstrated an improved toxicity as 1 μM in the presence of 4 μM copper completely blocked the growth of the pathogen (Fig. 5f). Increasing 8QN concentration to 2 μM in the presence of lower dose (2 μM) copper was also lethal to *S. parasitica* (Fig. S7).

### 3.3 BCS reverses the toxicity of ionophores-Cu^2+^ against *S. parasitica*

To further verify the role of copper in the observed toxicity against *S. parasitica* when combined with non-lethal doses of ionophores, we used the extracellular copper chelator bathocuproine sulfate disodium salt hydrate (BCS) to sequester copper in the extracellular growth media. The results confirmed that 10 μM TDD consistently inhibited *S. parasitica* growth in the 2222X culture media, and the addition of 8 μM copper highly enhanced the growth suppression (Fig. 6a). However, the addition of 250 μM BCS which prevented Cu^2+^ from entering the intracellular space (cytosol) reversed the inhibition induced by copper-ionophore (Fig. 6a). Increasing the copper concentration to 16 μM and 32 μM, respectively, in the presence of 10 μM TDD completely blocked *S. parasitica* growth. However, the lethal effect was reversed in the presence of 250 μM BCS. Meanwhile, the addition of 10 μM TDD and 250 μM BCS in the absence of copper was not lethal to *S. parasitica* which buttressed the role of copper in curving *S. parasitica* growth. A similar trend was observed for CLP and MPO (Fig. 6b and 6c). BCS exhibited the ability to reverse the lethal effect of CLP-Cu^2+^ and MPO-Cu^2+^ on *S. parasitica*. Remarkably, BCS in the absence of ionophore or copper has no lethal effect on the growth of *S. parasitica*. Therefore, it may not contribute to the observed modulatory activity of Cu^2+^ and ionophores against *S. parasitica* in this study. Surprisingly, CHI and DHQ in combination with 250 μM BCS resulted in total inhibition of *S. parasitica* growth (Fig. 6d and 6f). Addition of 8 μM copper to *S. parasitica* in the presence of 2 μM CHI supported the growth of the pathogen *in vitro*. However, the addition of BCS to the system resulted in the death of *S. parasitica* (Fig. 6d). We increased the concentration of copper to 16 μM in the presence of CHI and noticed that the growth of *S. parasitica* was slightly enhanced. More importantly, addition of BCS (250 μM) to the culture resulted in the death of *S. parasitica*. A similar pattern was recorded for DHQ (Fig. 6e), with copper leading to the improved growth rate of the pathogen, whereas the addition of BCS led to death. This suggests that CHI and DHQ likely share a common mechanism that differs from that of TDD, CLP, MPO and 8QN. Fig. 6f indeed showed that 8QN (8 μM) and 2 μM Cu^2+^ suppressed the growth of *S. parasitica* in comparable manner to TDD, CLP and MPO. Increasing the dose of Cu^2+^ to 4 μM in the presence of 8QN resulted in the death of the pathogen whereas addition of 250 μM BCS yielded a reversal of the lethal effect (Fig. 6f).

**Fig. 6.**
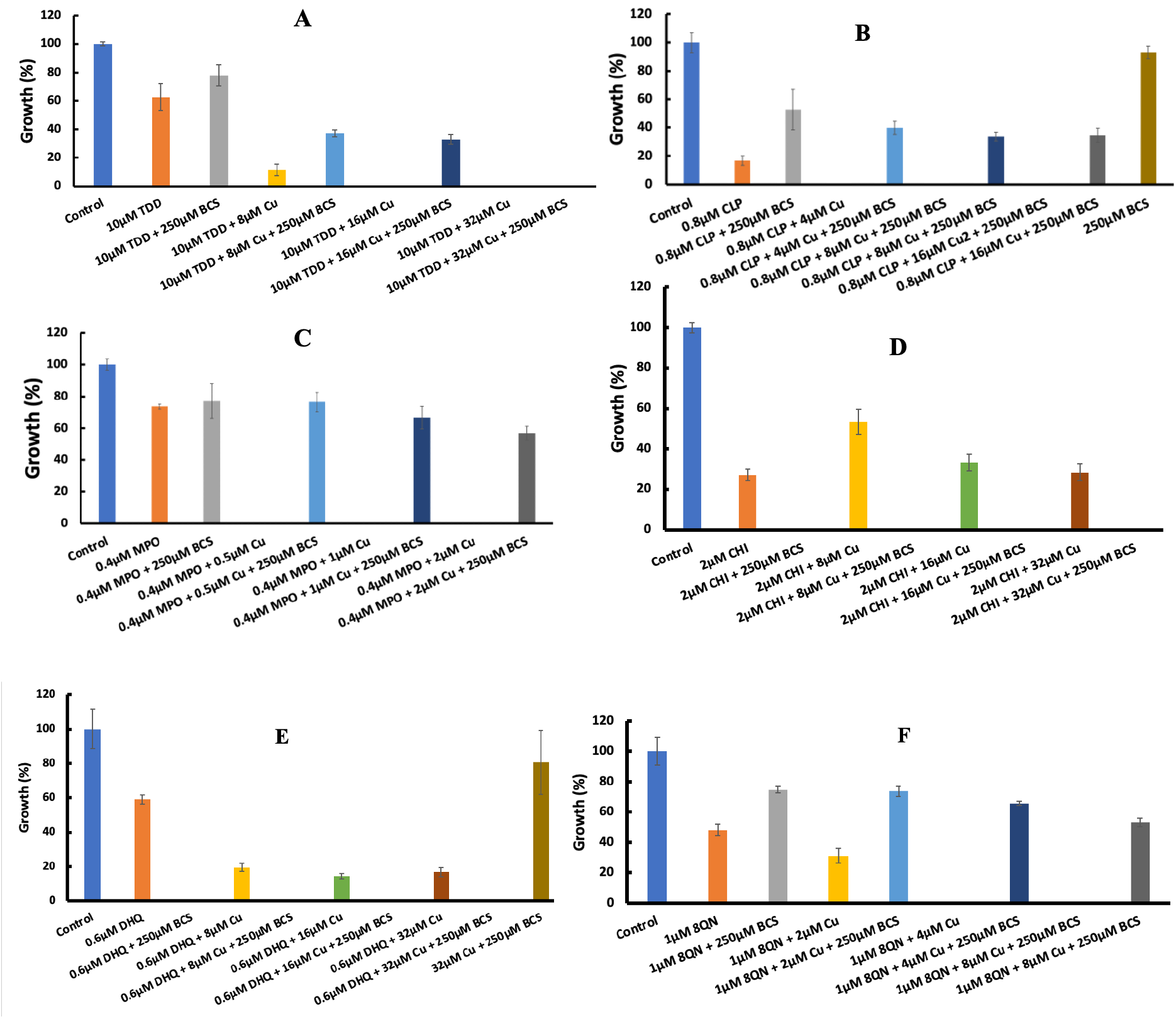
Effect of withholding Cu^2+^ in the extracellular space against *S. parasitica* growth *in vitro*. A combination of copper and ionophores was lethal to *S. parasitica* at low dose (32μM Cu^2+^). (a) 10 μM TDD, (b) 1 μM CLP, (c) 1 μM MPO, (d) 4 μM CHI, (e) 0.8 μM DHQ and (f) 8 μM 8QN. Growth was measured at 28°C after 5 days and normalized to the growth of control. Error bars represent the standard deviation of the mean of data from triplicate experiments.

### 3.4 Modulating *S. parasitica* growth on solid media with copper and ionophores

To mimic the effects of the Cu^2+^ - ionophore strategy on *S. parasitica* grown on the solid host, we prepared 2222X solid media plates and cultured the oomycete, treated with Cu^2+^ or ionophores for three days at room temperature. Data generated will corroborate the observed effect of copper and ionophores on *S. parasitica* carried out in 2222X media (solution). We determined the dose-dependent effect of copper (Fig. S8), ionophores (Fig. 7) and Cu^2+^-ionophore (Fig. 8) on the mycelial growth of *S. parasitica*. The inhibitory effect of each ionophore against the pathogen was dose-dependent. However, a relatively higher dose was required to completely block the mycelial growth of *S. parasitica* on the solid media than in solution. Reason?

**Fig. 7.**
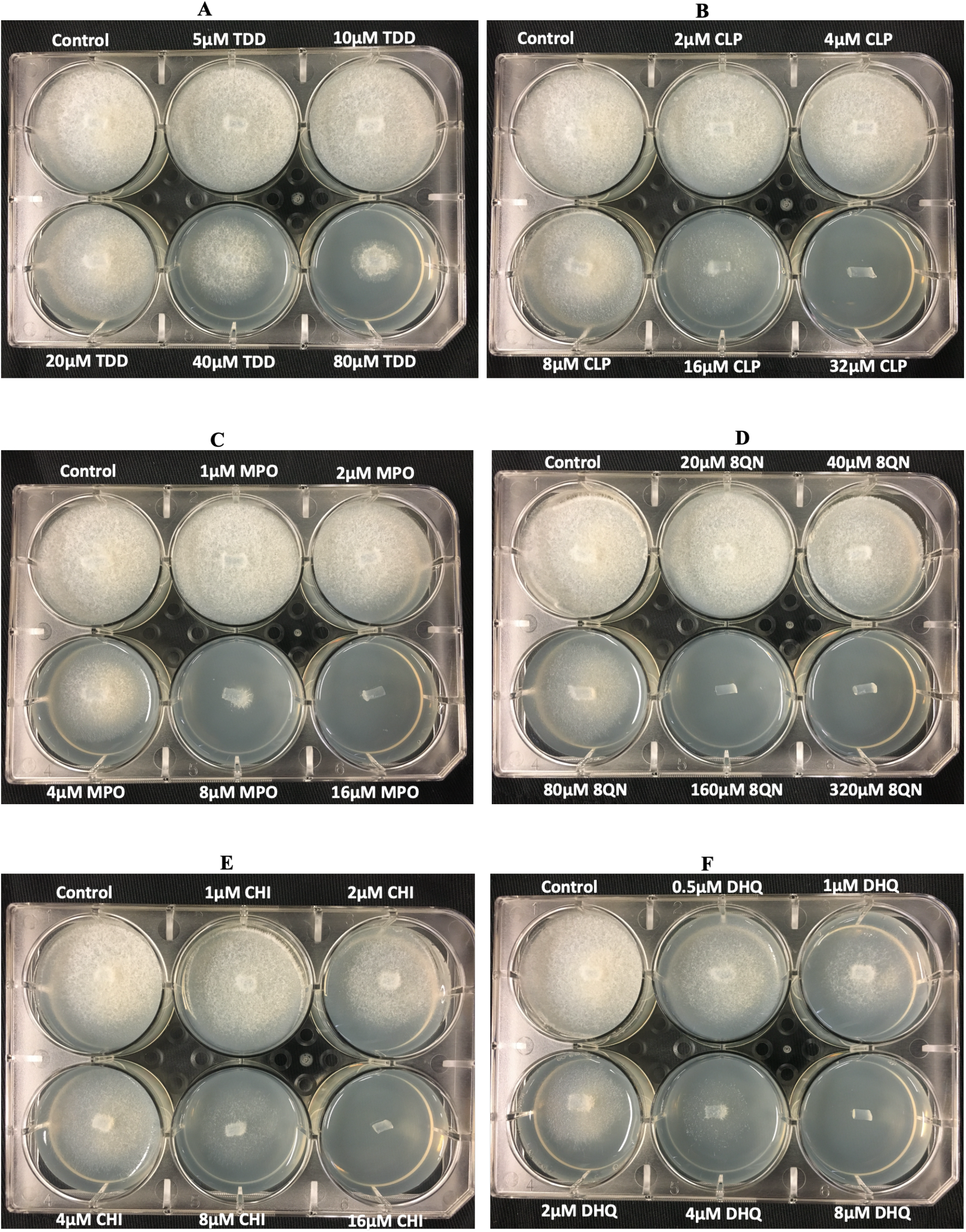
Effect of ionophores on *S. parasitica* grown on solid media. The effect of each ionophore is dose dependent. (a) TDD (0 μM – 80 μM), (b) CLP (0 μM – 32 μM), (c) MPO (0 μM – 16 μM), (d) 8QN (0 μM – 320 μM), (e) CHI (0 μM – 16 μM) and (f) DHQ (0 μM – 8 μM). Images were taken after 3 days at room temperature to determine growth of *S. parasitica* mycelia.

### 3.5 Mechanisms underlying the toxicity of Cu^2+^-ionophore against *S. parasitica*

To verify whether metal-mediated oxidative stress plays a direct role in the copper and ionophore toxicity observed in this study, we utilized the oxyblot kit to derivatize and detect oxidized proteins in *S. parasitica* lysates under various treatment conditions (Fig.10). Similar levels of oxidized proteins are observed across 2-hour acute treatment conditions involving combinations of MPO, Cu and BCS (Fig 10A). This suggests that the mechanisms underlying the Cu^2+^ - ionophore toxicity in *S. parasitica* may be targeted and not involve a general increase of metal-mediated oxidative stress. How?

## 4. DISCUSSION

Research efforts have continued till date to seek alternatives and develop new strategies for the control of Saprolegniasis in aquaculture, with special attention on toxicity and safety (Ali et al., 2019; Meneses et al., 2022; Werner et al., 2020; Zhang et al., 2019). Based on the known antimicrobial activity of copper, we evaluated a new strategy that can increase the antimicrobial efficiency of copper against *S. parasitica* by combining low doses of ionophores and copper sulfate (Straus et al., 2020). Therefore, we prepared a synthetically defined media (2222X media) that can successfully and reproducibly support *S. parasitica* growth. In this 2222X growth media, 100 μM Cu sulfate can reproducibly suppress 50% of total *S. parasitica* mycelium growth (Fig. 2). This copper concentration is consistent with other reports using copper sulfate as an anti-*Saprolegnia* treatment to suppress *S. parasitica* growth on largemouth bass and catfish eggs (Straus et al., 2020; Straus, Mitchell, Carter, et al., 2009). Screening eighteen potential Cu-binding small molecules revealed six compounds that were able to completely suppress *S. parasitica* growth at a concentration of 10 μM in combination with 100 μM Cu-Sulfate (Fig 3.). Among these six promising compounds, four displayed Cu-dependent antimicrobial activity, and two inhibited *S. parasitica* growth regardless of exogenous copper concentration.

Further investigation into the effectiveness of these six chemical compounds was conducted to determine their IC_50_ doses in 2222X growth media alone. Interestingly, all six compounds were effective at inhibiting Saprolegnia growth with lethal doses in the micromolar range (Fig. 4). The compound DHQ was the most effective with a lethal dose at 0.8 μM (Fig. 4E), followed by MPO and CLP with a lethal dose of approximately 1 μM. The compounds CHI and 8QN were slightly less effective, with a lethal dose accruing at 4 μM and 8 μM, respectively. In the absence of Cu, a higher 20 μM concentration of TDD was needed. These data contribute to the ongoing research efforts to fully understand the bioactivities of ionophores, including their cellular toxicity, antimicrobial and antioxidant effects (Song et al., 2019). Indeed, these findings are comparable to the reported effect of ionophores against the fungal pathogen *C. neoformans* (Helsel et al., 2017).

We next investigated the effectiveness of the six chemical compounds identified in our initial screen to increase Cu toxicity on *S. parasitica*. Using a dose of each compound at or below the IC_50_, we systematically altered supplemental copper concentrations to determine the minimal copper dose needed to completely suppress *S. parasitica* propagation. We observed that increasing the copper dose at the fixed ionophore concentration revealed two distinct trends in *S. parasitica* Cu-dependent growth behavior (Fig. 5). The chemicals TDD, CLP, MPO, and 8QN, which are more active with increasing Cu concentrations, were considered as one group (Fig 5. A.B.C.F). However, DHQ and CHI, which display moderate growth inhibition at both low and high levels of supplemental copper were considered a second defined group. The anti-saprolegnia activity of DHQ and CHI is abolished at a copper concentration of 1 μM but then restored at higher levels of supplemental copper. These trends in *S. parasitca* copper-dependent growth behavior are largely conserved when applied to *S. parasitica* solid-supported growth assays (Fig. 8). The ability to inhibit *S. parasitica*, both in liquid culture as well as solid-supported growth formats, suggests that the combination of copper sulfate and small chemical agents may effectively treat *S. parasitica* infection on both developing fish embryos as well as adult fish.

**Fig. 8.**
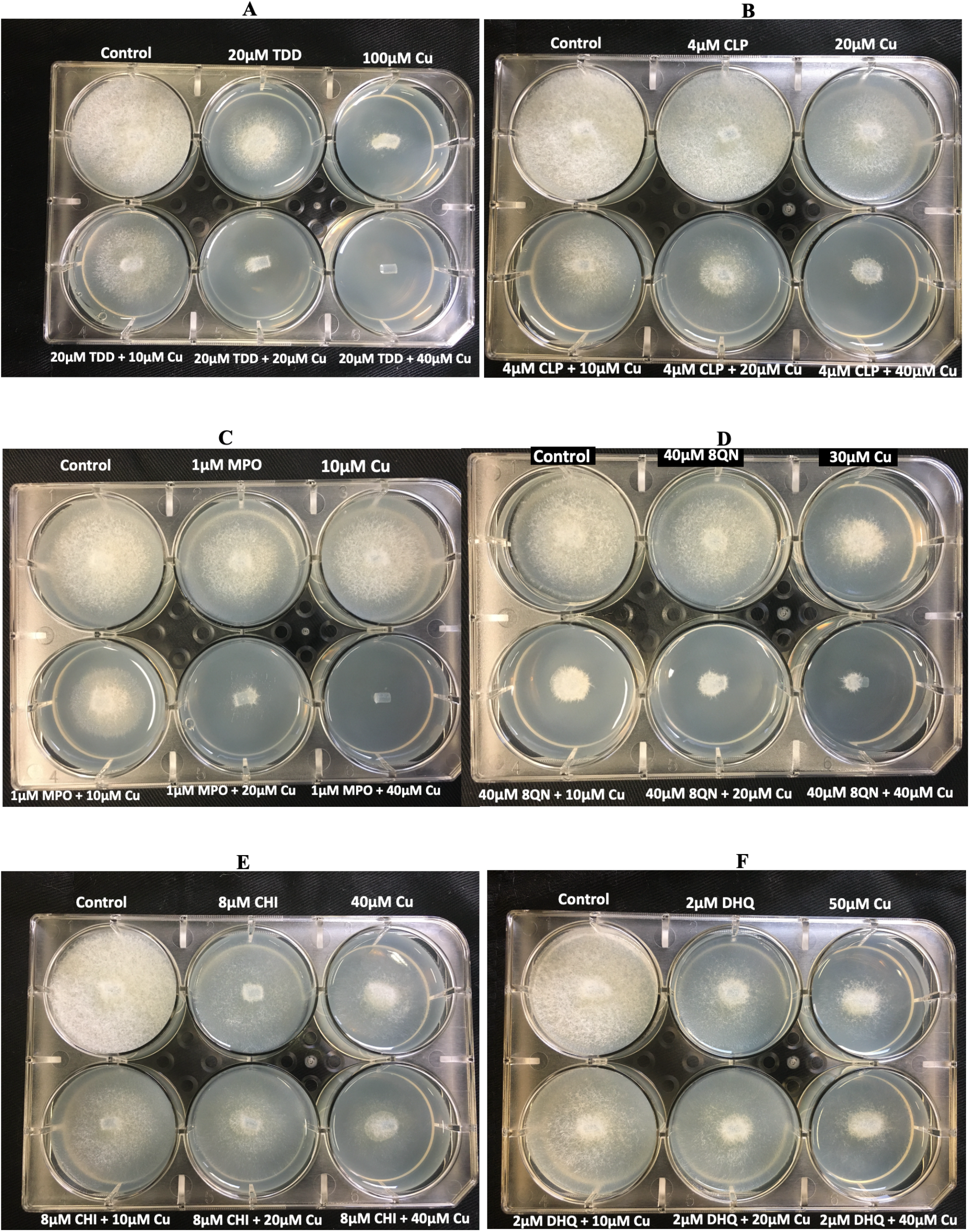
Effect of Cu^2+^-ionophore on *S. parasitica* grown on solid media. The effect of each ionophore is dose-dependent. (a) TDD (0 μM – 80 μM), (b) CLP (0 μM – 32 μM), (c) MPO (0 μM – 16 μM), (d) 8QN (0 μM – 320 μM), (e) CHI (0 μM – 16 μM) and (f) DHQ (0 μM – 8 μM). Images were taken after 3 days at room temperature to determine growth of *S. parasitica* mycelia (Tedesco et al., 2019).

The cell impermeable copper-specific chelator BCS was used to restrict supplemental copper to the extracellular environment to clarify copper’s role in the observed anti-*Saprolegnia* activity. The addition of BCS can completely reverse the lethality of the group one chemicals (i.e. TDD, MPO, CLP and 8QN) while increasing the effectiveness of the group two chemicals (i.e. DHQ and CHI; (Fig. 6)). Based on these experiments, we propose the mode of action for group one chemicals require copper to suppress *S. parasitica* growth whereas group two chemicals operate in a copper independent mechanism. This suggests that group one chemicals function as Cu-ionophores (Fig. 9a,b) to disrupt cellular copper homeostasis by trafficking copper ions across the plasma membrane that does not rely on metal ion transports including the divalent metal transporter (DMT) or Copper-transporters (CTR) families (Lingli & Lin, 2013; Mandal et al., 2020).

**Fig. 9.**
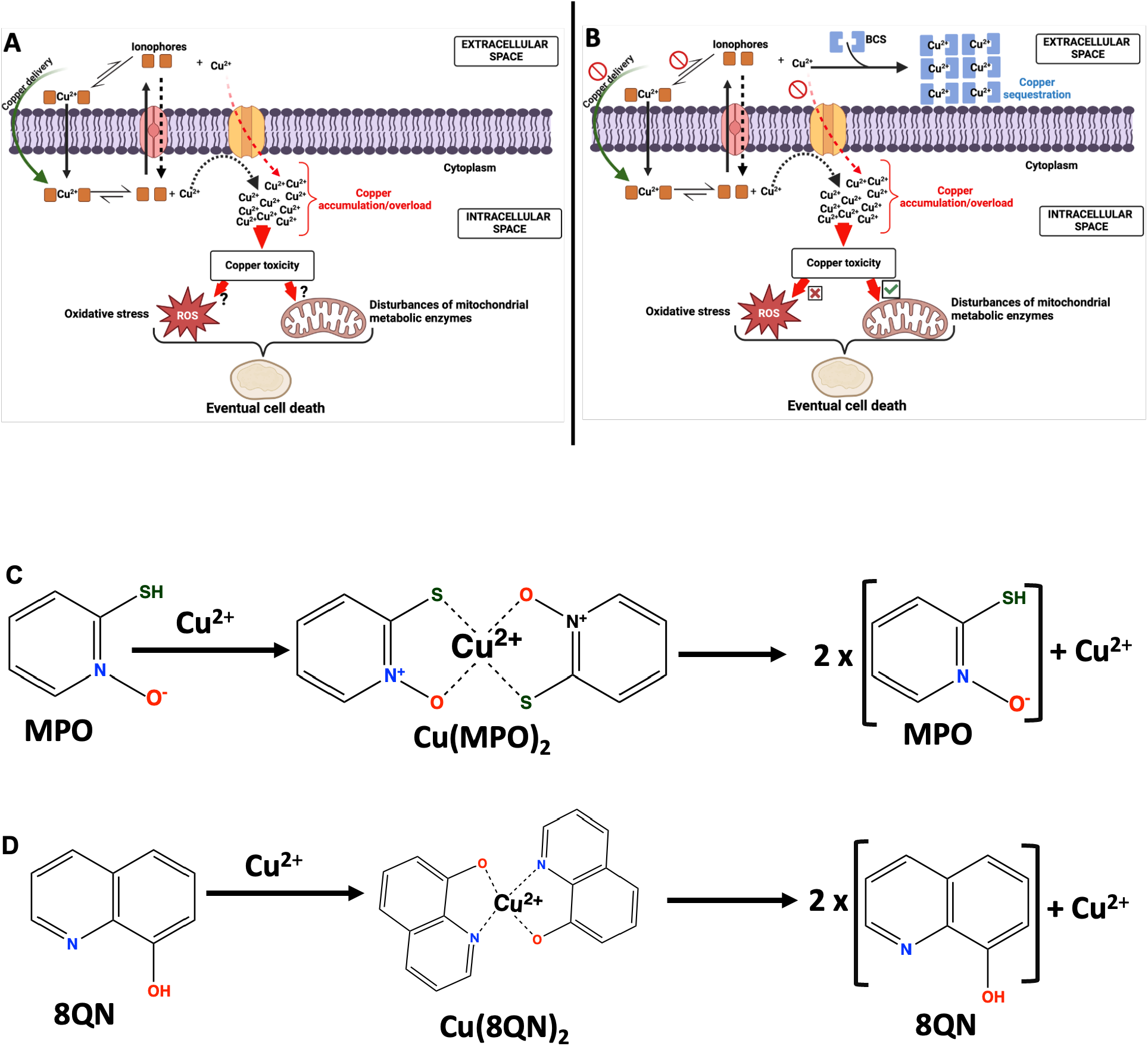
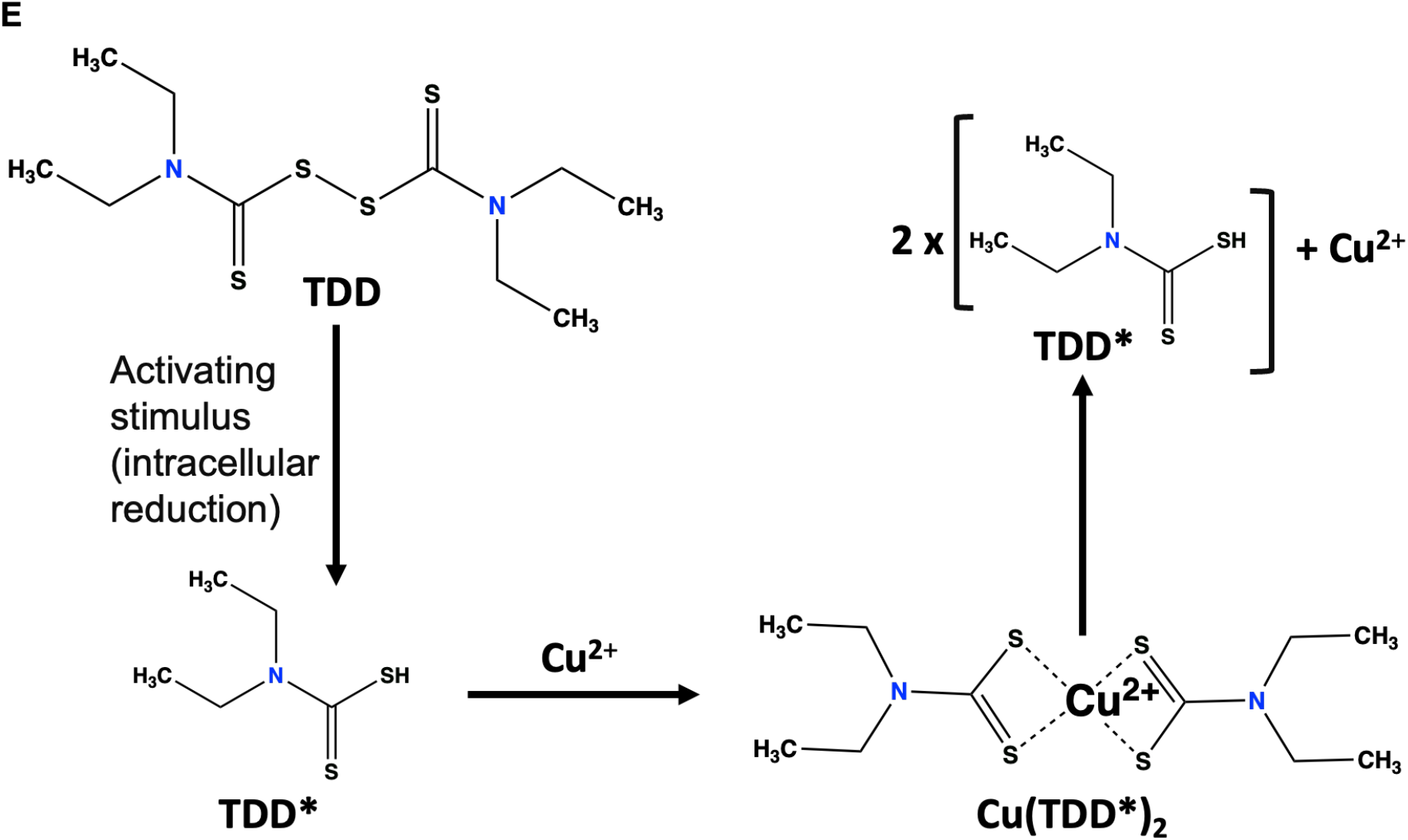
Schematic description of molecular mechanism underlying copper-ionophore inhibition *S. parasitica* growth. (a) Depicts mechanism of TDD, CLP, MPO and 8QN in the presence of copper. Copper is delivered in complex with the ionophores (ionophore - Cu^2+^) to *S. parasitica* cytoplasm, resulting in the over-accumulation of the metal with its associated toxicity. (b) Addition of BCS rescued the growth of *S. parasitica* by keeping copper in the extracellular space. BCS competes with the ionophores and favorably bind copper and sequestrating the metal in the extracellular space. The complex formed between copper and (c) MPO, (d) 8QN, (e) TDD which requires an activating stimulus prior copper-binding.

Group one chemicals, such as MPO and its’ respective metal salts, have been shown to interact with the cell membrane to transport metal ions into the cytoplasm, resulting in metal accumulation, which may alter the metabolic processes in the cell (Reeder et al., 2011). Two MPO molecules can bind a single divalent copper ion to form Cu(MPO)_2_ (Mishra et al., 2023), as shown in Figure 9c-e. We hypothesize this neutral Cu(MPO)_2_ complex is responsible for cytosolic Cu-delivery. Interestingly, MPO has successfully been used to combat fungal pathogens such as *Malassezia restricta and Malassezia glabosa* in human scalps (Park et al., 2018). Similarly, its zinc salt (MPOZ) is applied in commercial skin care products to treat psoriasis, seborrheic dermatitis, eczema, and acne (Mangion et al., 2021; Sadeghian et al., 2011). Similarly, the compounds 8QN, TDD, and CLP also have the capacity to bind metal ions and exhibit broad-spectrum antimicrobial activities (Abrams et al., 1991; Helsel et al., 2017). Our data suggest that the 8QN, TDD, and CLP modes of action for *S. parasitica* growth inhibition may involve metal binding and trafficking, specifically with copper ions (Helsel & Franz, 2015; Mishra et al., 2023).

To test whether the mode of action of Cu^2+^-ionophore treatment on *S. parasitica* involved a mechanism involving metal-mediated oxidative stress, we performed a time-course experiment to quantify cellular protein oxidation. Using the OxyBlot immunodetection assay on total cell lysates revealed no significant difference in oxidative protein levels among MPO-Cu treated samples relative to control conditions (Fig. 10). This suggests that MPO-Cu toxicity may not involve global changes in cellular oxidative stress but rather a more specific pathway. Excess copper has recently been shown to alter mitochondrial metabolic enzymes and related protein-binding processes (Kahlson & Dixon, 2022). We suspect that this alteration, termed cuproptosis, may play a role in the observed Cu-ionophore toxicity in the current study. This pathway will be further explored in our future project.

**Fig. 10.**
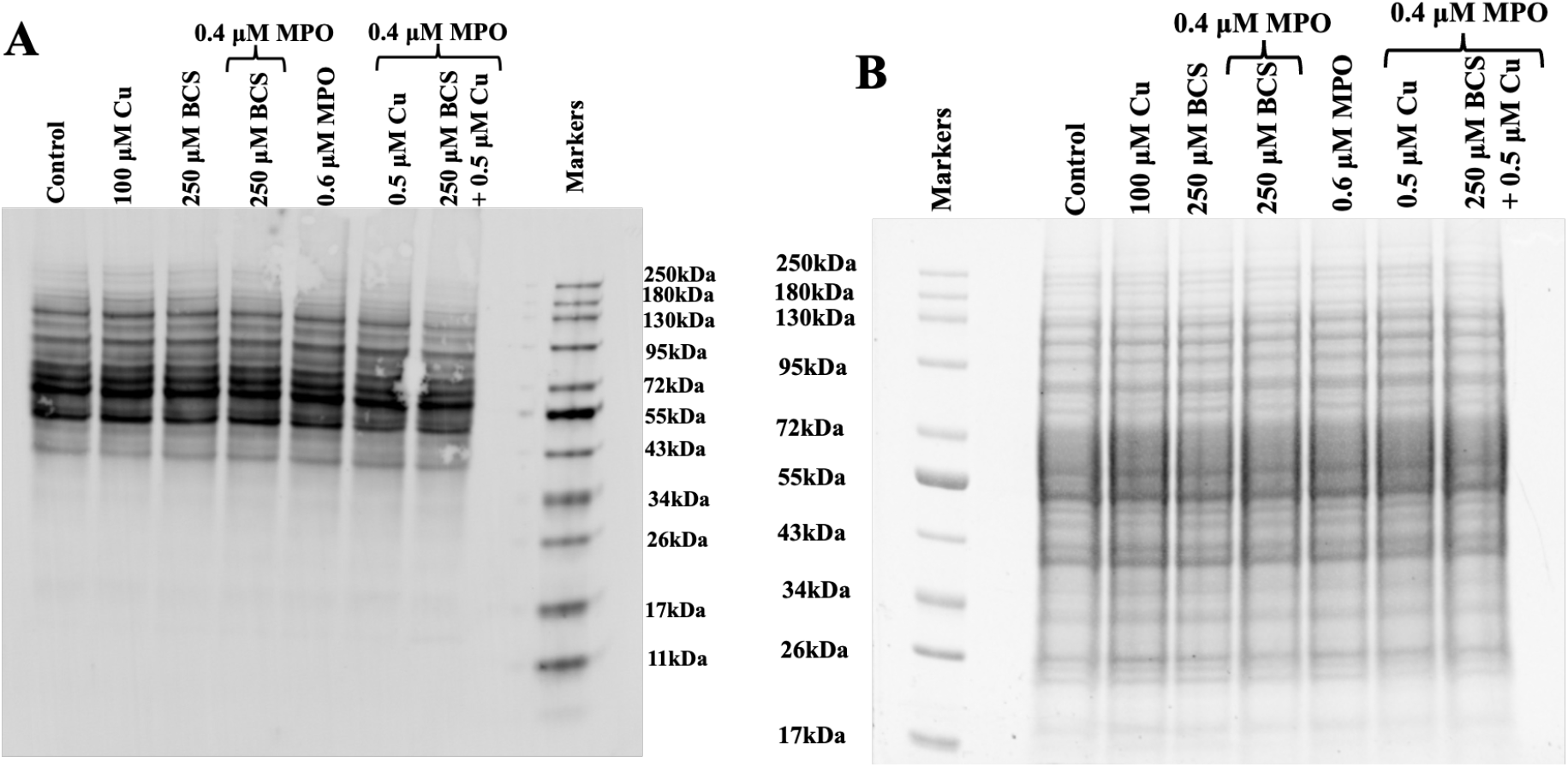
Mechanism underlying the ionophore-copper toxicity in *S. parasitica*. (a) Determining the possible contribution of oxidative stress in the ionophore (MPO)-copper toxicity against *S. parasitica*, the carbonyl groups in 80 μg *S. parasitica* protein samples were derivatized to 2,4-dinitrophenylhydrazone (DNP) in the presence of 6% sodium dodecyl sulfate (SDS), followed by an incubation at 25°C for 15 minutes. The reaction was stopped by adding the neutralization solution and the samples run on a 12% SDS-PAGE. (b) Total proteins in *S. parasitica* on an SDS-PAGE gel. Consistent protein bands were observed in the pathogen under all treatments.

## 5. CONCLUSION

The oomycete pathogen *S. parasitica* can cause large production losses across the entire lifespan of catfish and salmon aquaculture production stages from hatcheries to harvest. Herein, we tested and developed a new strategy for boosting copper toxicity against the oomycete by combining ionophores with minimal doses of copper sulfate. The six chemical agents identified in this study were effective at inhibiting *S. parasitica* at low micromolar concentrations in the presence of copper. This observation was verified using a copper-specific and cell impermeable chelator (BCS) to restrict bioavailable cellular copper level, reversing the toxicity and growth of the pathogen. This suggests that the growth of *S. parasitica* can be effectively controlled by using a low dose of copper and ionophore. Our laboratory’s efforts are ongoing to assess the effectiveness of Cu-ionophore combinations in inhibiting *S. parasitica* growth on developing catfish embryos.

## ACKNOWLEDGEMENTS

We would also like to thank Dr. Mike Mathews, Dr. John Hawke, and Chris Darnall, and the members of the A.E. Wood Fish Hatchery for supports, helpful advice and discussions.

## AUTHOR CONTRIBUTIONS

**Tomisin Happy Ogunwa:** Investigation, analyses, writing, proofreading

**Madison Grace Thornhill:** Methodology, validation, investigation

**Daniel Ledezma:** Methodology, investigation, preliminary data collection

**Ryan Loren Peterson:** Funding acquisition, project design and supervision, review, editing.

