## Supplemental files for "Modulation of *Saprolegnia parasitica* growth with copper and ionophores"

**Running title: Cu/ionophores inhibit Saprolegniasis**

Tomisin Happy Ogunwa, Madison Grace Thornhill, Daniel Ledezma and Ryan Loren Peterson†
Department of Chemistry, Texas State University, 601 University Drive, San Marcos, Texas, United States, 78666.
† Corresponding author

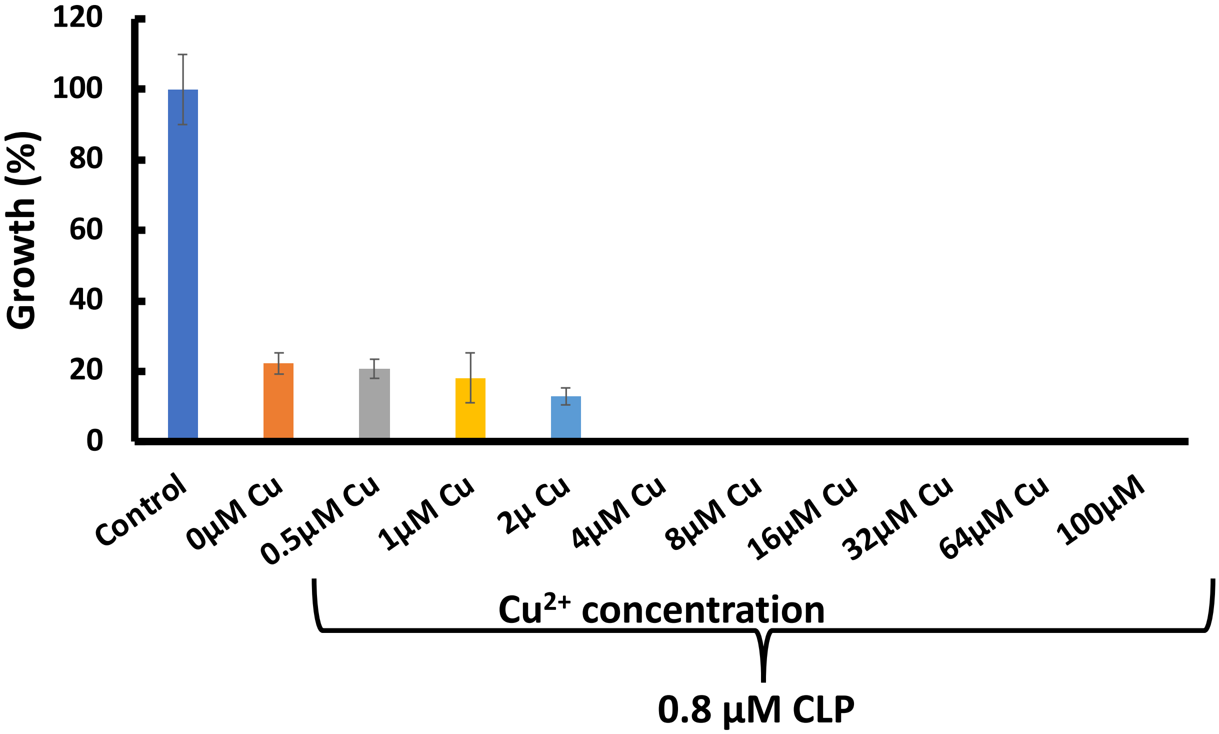


**Fig. S1.** Effect of copper - CLP (0.8 µM) treatment on *S. parasitica* growth *in vitro*. Total suppression of growth in *S. parasitica* was observed at 4 µM copper in the presence of 0.8 µM CLP.

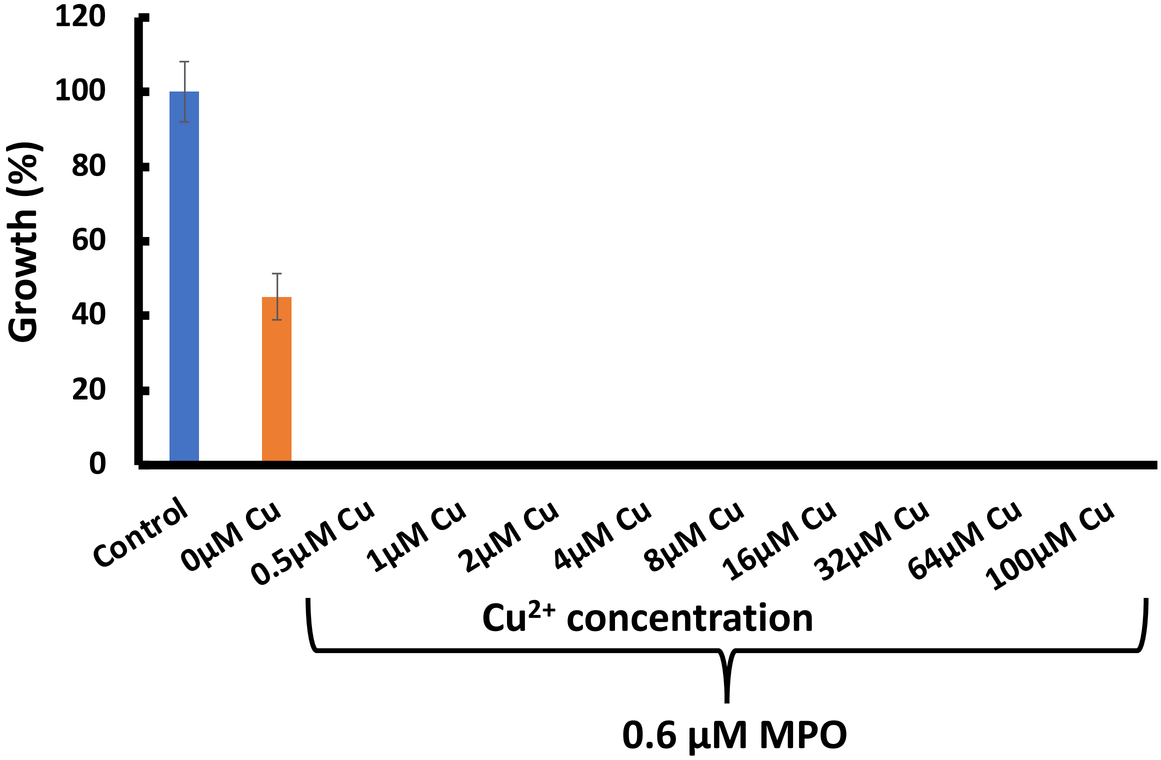


**Fig. S2**. Effect of copper – MPO (0.6 µM) treatment on *S. parasitica* growth *in vitro*. Total suppression of growth in *S. parasitica* was observed at 0.5 µM copper in the presence of 0.6 µM MPO.

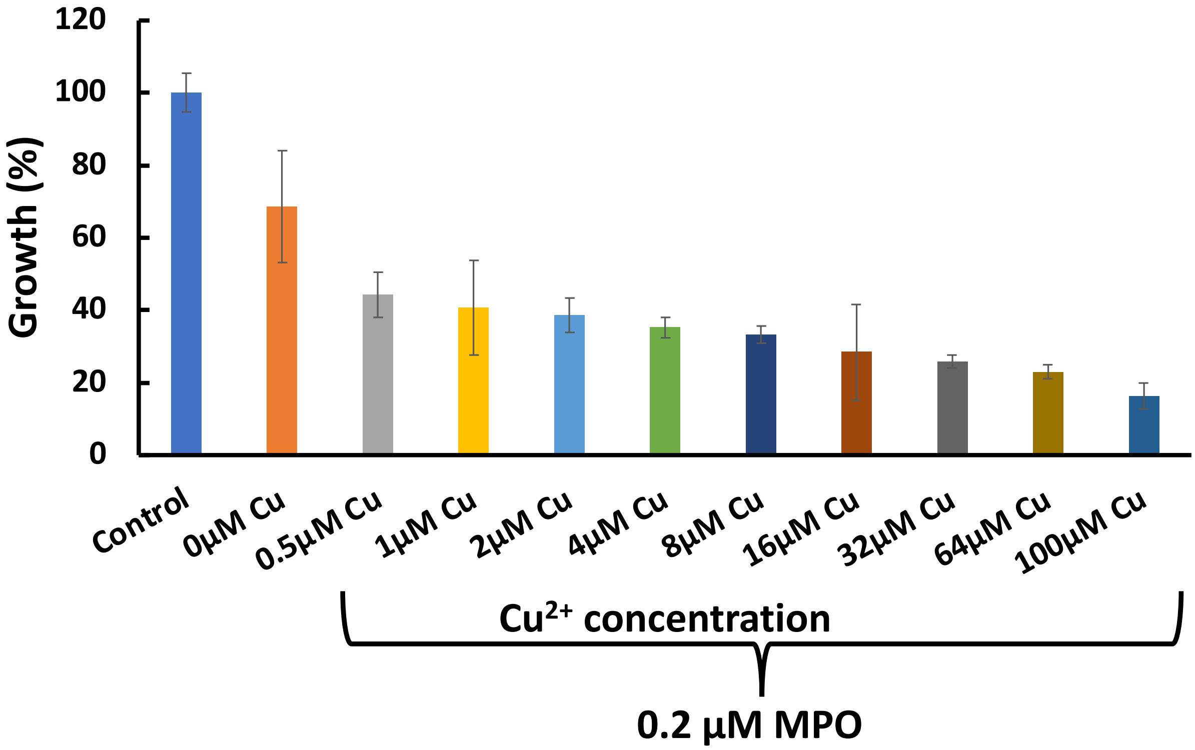


**Fig. S3.** Effect of copper – MPO (0.2 µM) treatment on *S. parasitica* growth *in vitro*. *S. parasitica* survived in the presence of 100 µM copper and 0.2 µM MPO.

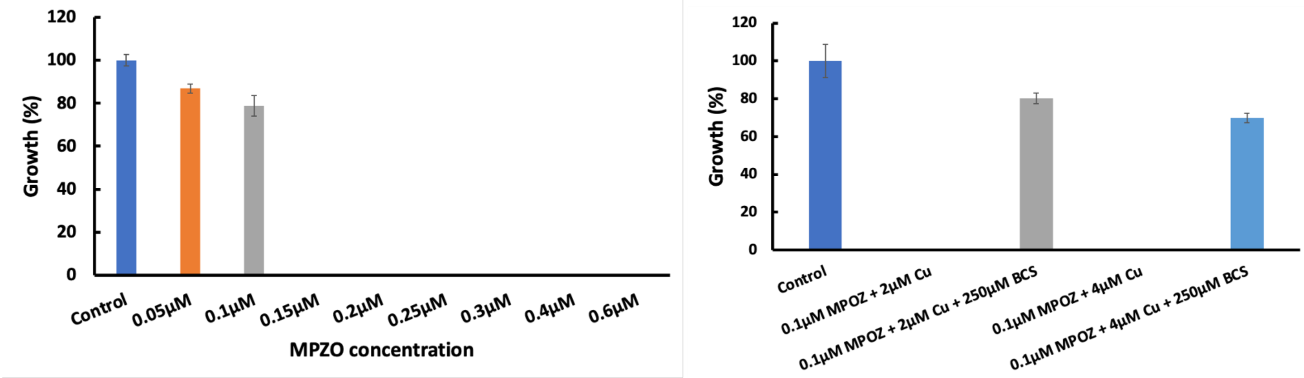


**Fig. S4**. (A) Dose-dependent assay of MPOZ against *S. parasitica*. Lethal dose for MPOZ was 0.15 µM as determined *in vitro*. (B) Treatment of *S. parasitica* with copper - MPOZ and copper - MPOZ – BCS *in vitro*. Total suppression of growth in *S. parasitica* was observed in the presence of 0.1 µM MPOZ combined with 2 µM or 4 µM copper. This effect was reversed by the addition of BCS (250 µM).

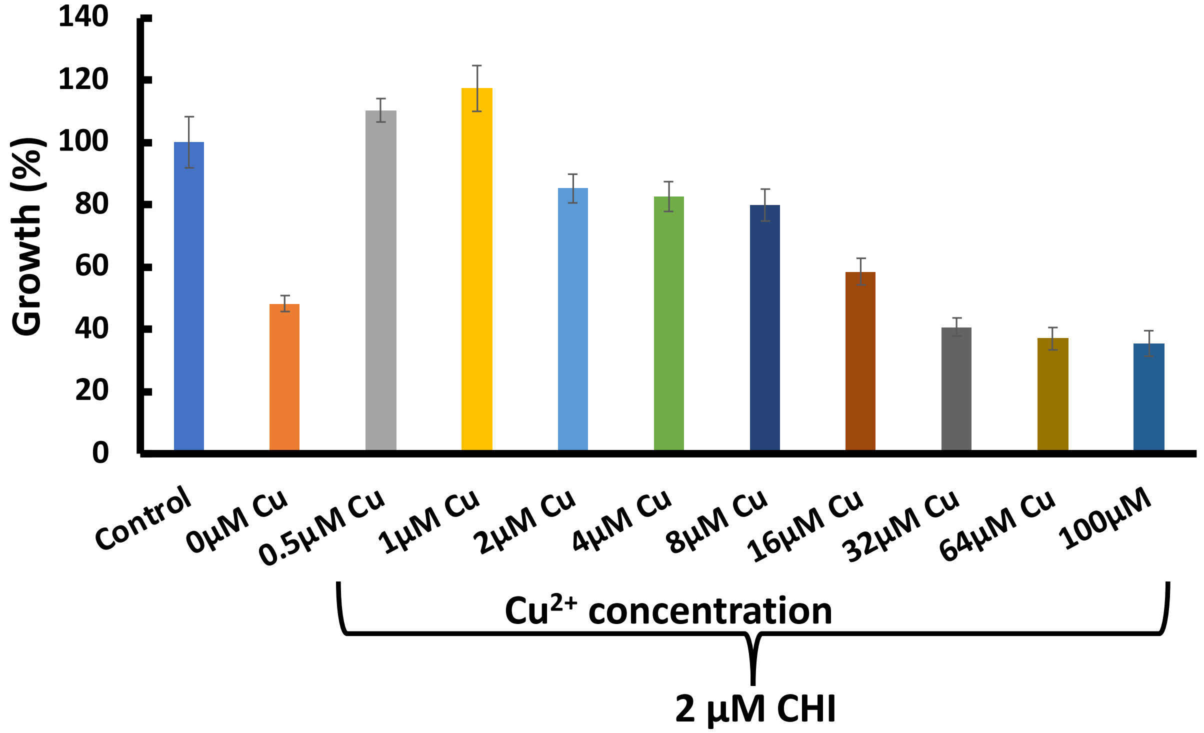


**Fig. S5.** Effect of copper – CHI (2 µM) on *S. parasitica* growth *in vitro*. An enhanced growth of *S. parasitica* was seen at 2 µM CHI dose and lower doses of copper (0.5 µM – 1µM). A dose-dependent suppression of *S. parasitica* growth was thereafter observed at higher doses of copper.

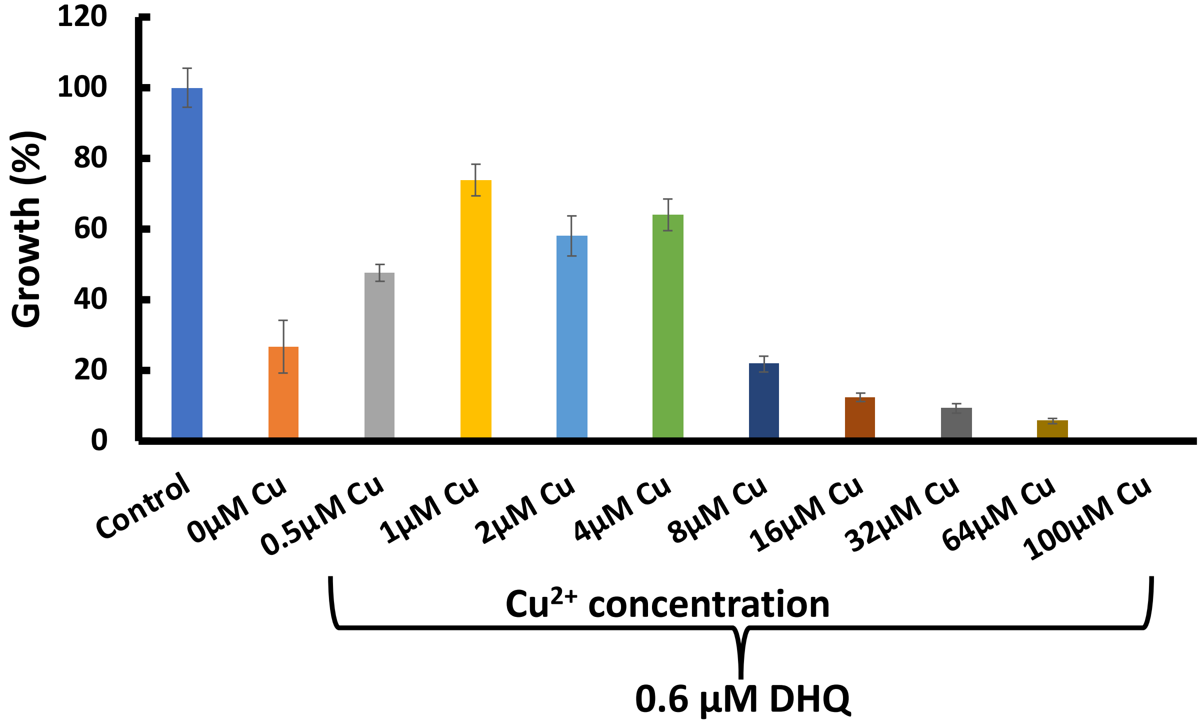

**Fig. S6**. Effect of copper – DHQ (0.6 µM) on *S. parasitica* growth *in vitro*. An enhanced growth of *S. parasitica* was observed at 0.6 µM DHQ concentration and 1 µM copper.

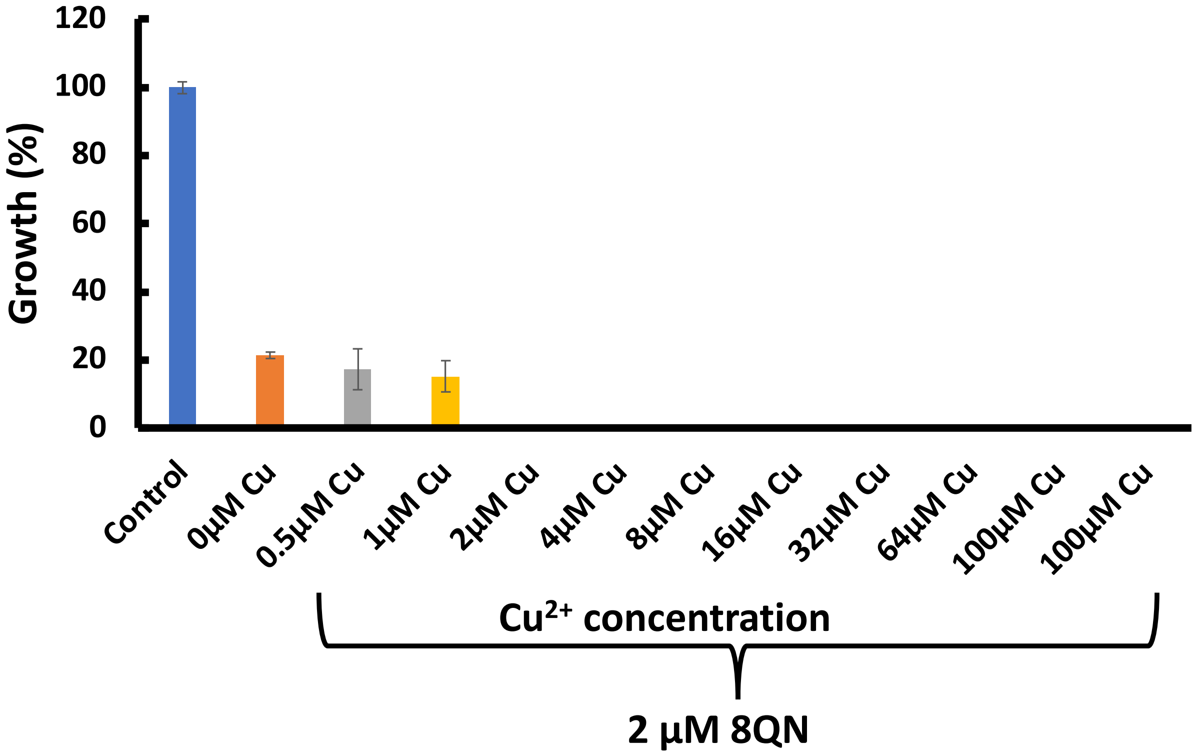


**Fig. S7**. Effect of copper – 8QN (2 µM) on *S. parasitica* growth *in vitro*. A combination of 2 µM 8QN and 2 µM copper was lethal to *S. parasitica*.

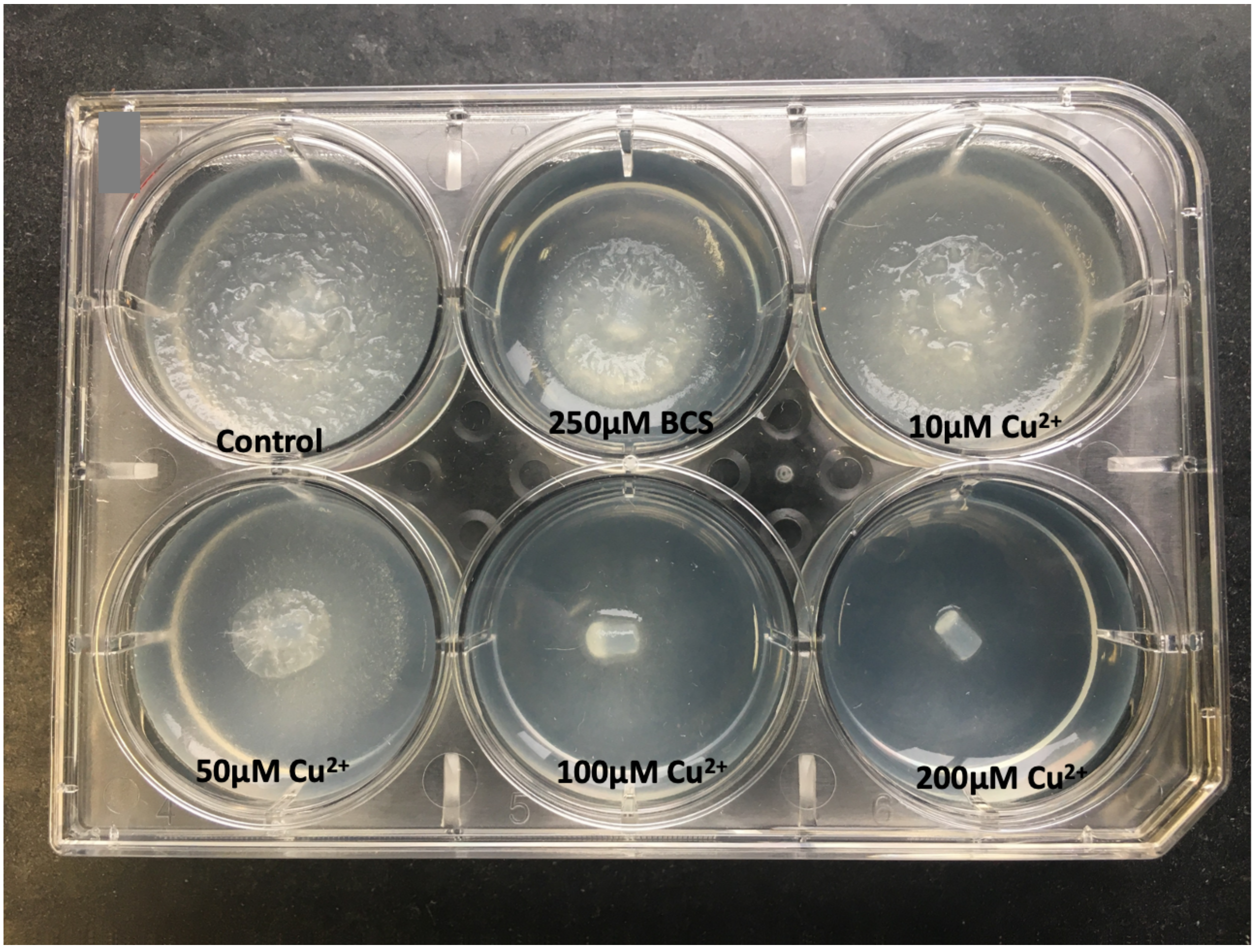

**Fig. S8**. Effect of varying copper concentration on the growth of *S. parasitica* on solid media in a six-well plate. Copper curved *S. parasitica* growth in a dose-dependent manner.

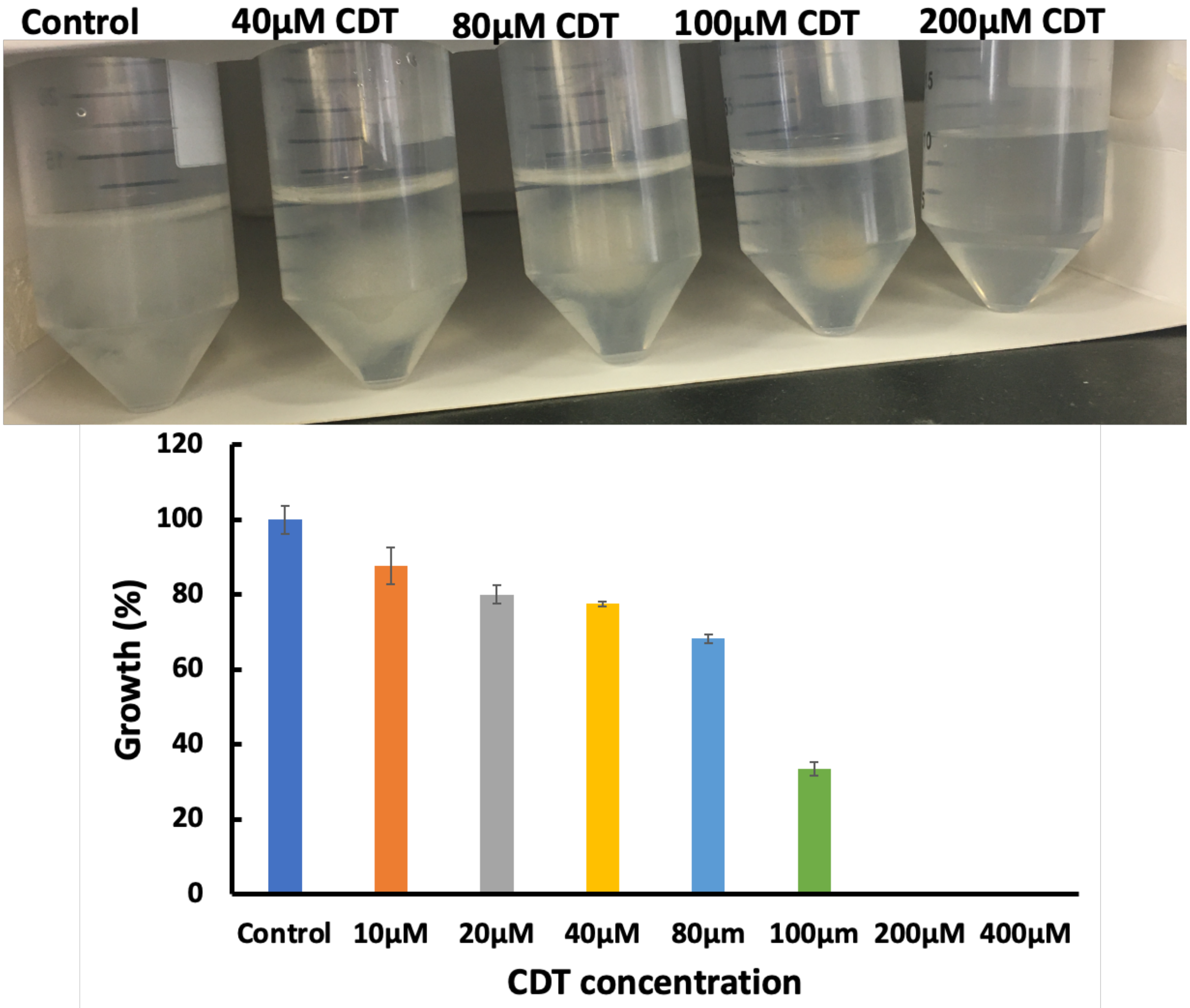


**Fig. S9**. Dose dependent effect of CDT on the growth of *S. parasitica in vitro*. $IC_{50}$ was 100 µM whereas lethal dose 200 µM.
